# *Drosophila* Wg and Evi/Wntless dissociation occurs post apical internalization in the late endosomes

**DOI:** 10.1101/2023.12.07.570516

**Authors:** Satyam Sharma, Varun Chaudhary

## Abstract

Signaling pathways activated by secreted Wnt ligands play an essential role in tissue development and the progression of diseases, like cancer. Secretion of the lipid-modified Wnt proteins is tightly regulated by a repertoire of intracellular factors. For instance, a membrane protein, Evi/Wntless, interacts with the Wnt ligand in the ER, and it is essential for its further trafficking and release in the extracellular space. After dissociating from the Wnt, the Wnt-unbound Evi is recycled back to the ER via Golgi. However, where in this trafficking path Wnt proteins dissociate from Evi remains unclear. Here, we have used the *Drosophila* wing epithelium to trace the route of the Evi-Wg (Wnt homolog) complex leading up to their separation. In these polarized cells, Wg is first trafficked to the apical surface, however, the secretion of Wg is believed to occur post-internalization via recycling. Our results show that the Evi-Wg complex is internalized from the apical surface. Furthermore, using antibodies that specifically label the Wnt-unbound Evi, we show that Evi and Wg separation occurs post-internalization in the acidic endosomes. These results refine our understanding of the polarized trafficking of Wg and highlight the importance of Wg endocytosis in its secondary secretion.

## INTRODUCTION

Wnts are highly conserved secreted ligands that can activate signaling pathways essential for various cellular and developmental processes, for example, cell proliferation, cell fate specification, and tissue patterning. For a precise regulation of these processes, secretion of Wnt ligands occurs in a tightly controlled manner. It initiates with their post-translational modification via glycosylation in the endoplasmic reticulum (ER) followed by palmitoylation by a membrane-bound ER-resident O-acyltransferase Porcupine (Porc) (Kadowaki et al., 1996; Komekado et al., 2007; Kurayoshi et al., 2007; Takada et al., 2006; Van den Heuvel et al., 1993; Willert et al., 2003). With some exceptions, these modifications facilitate further transport of Wnt proteins via a dedicated eight-pass transmembrane cargo-receptor protein Evenness interrupted (Evi; also known as Wntless, Sprinter, MIG-14 and GPR177) (Bänziger et al., 2006; Bartscherer et al., 2006; Goodman et al., 2006; Herr and Basler, 2012). Structural analysis of Evi and Wnt interactions suggested that Wnt is transferred directly from Porc to Evi in the ER, and palmitoylation of Wnt allows its interaction with the hydrophobic pockets in the extracellular domain of Evi (Nygaard et al., 2021; Zhong et al., 2021). The Wnt-Evi complex is further transferred from ER to the Golgi with the help of p24 family proteins in the COPII-coated vesicles and then to the plasma membrane via Rab8a-dependent trafficking (Buechling et al., 2011; Castillon et al., 2011; Das et al., 2015; Port et al., 2011). Following separation from the Wnt, the Wnt-unbound Evi (‘free’ Evi) is recycled back to the Golgi in a retromer-dependent process (Belenkaya et al., 2008; Franch-Marro et al., 2008; Kim et al., 2009; Pan et al., 2008; Port et al., 2008; Yang et al., 2008) and then to the ER via COPI vesicles (Yu et al., 2014). This step protects Evi from lysosomal degradation and facilitates further secretion of Wnt proteins (Coudreuse et al., 2006; Prasad and Clark, 2006).

Studies on Wnt protein secretion using the developing *Drosophila* larval wing epithelium have played an important role in the identification of several essential factors in the Wnt secretory pathway. In this polarized wing epithelium, Wingless (Wg; a homolog of mammalian Wnt1) is produced by a narrow stripe of cells along the dorsal-ventral (DV) boundary. In the producing cells, intracellular Wg, as well as Evi, are present at higher levels towards the apical side (Port et al., 2008; Simmonds et al., 2001; Yamazaki et al., 2016). However, Wg is released in the extracellular space from both the apical and basolateral sides of the producing cells. The extracellular dispersion of Wg creates a symmetric concentration gradient on both sides of the DV boundary, which activates differential gene expression in a concentration-dependent manner (Strigini and Cohen, 2000; Zecca et al., 1996). However, the apical pool of Wg is believed to be functionally more active (Chaudhary and Boutros, 2019).

Wg is also shown to be internalized by the producing cells from the apical surface, which brings Wg to the endosomal compartments (Marois et al., 2006; Pfeiffer et al., 2002). Following internalization, multiple downstream routes for Wg trafficking have been suggested; for example, 1) Wg can be recycled back to the apical surface with the help of Rab4 and Ykt6 (Linnemannstöns et al., 2020). 2) Wg has been shown to undergo apical to basolateral transcytosis with the help of Dlp, Godzilla, and Klp98A (Gallet et al., 2008; Witte et al., 2020; Yamazaki et al., 2016). 3) Internalized Wg and other mammalian Wnts are trafficked to the multivesicular bodies (MVB), where they are packaged on the intraluminal vesicles, which are released as exosomes (Beckett et al., 2013; Gross et al., 2012; Korkut et al., 2009).

So far, studies have largely focused on tracking the intracellular route of Wnt/Wg protein, while the intracellular trafficking route of the Evi-Wg complex leading up to their separation remains largely unknown. Studies using cultured human cells have suggested that the separation of Wnt3a and Evi proteins requires acidic pH in the vesicles (Coombs et al., 2010). However, whether this occurs in the anterograde route or if Wnt and Evi are internalized together and the separation occurs post-internalization remains unclear.

Here, we have traced the route of the Evi-Wg complex in the polarized wing epithelial cells. We generated antibodies targeting the extracellular domain of Evi, which specifically detected the Wnt-unbound or ‘free’ Evi. This allowed us to probe the site for the separation of Wg and Evi. Our results show that the Evi-Wg complex reaches the apical membrane, which is followed by their co-internalization. Furthermore, we find that Evi and Wg separation occurs in the acidic late endosomes, following their apical internalization. These results suggest that apart from apical recycling and apical to basolateral transcytosis, internalization of Wg could also facilitate its dissociation from the cargo-receptor Evi.

## RESULTS

### Evi remains bound to Wg on the apical surface of the columnar cells

To understand the trafficking route of the Evi-Wg complex in the polarized wing epithelial cells, we first asked if Evi is transported to the cell membrane in the producing cells. To this end, we generated an antibody targeting the first and also the largest extracellular loop (α-Evi-ECD) (see materials and methods). In parallel, we also regenerated the published Evi antibody targeting the intracellular C-term domain (α-Evi-CTD) (Port et al., 2008). Next, we analyzed the specificity of the Evi-CTD and Evi-ECD antibodies using RNAi-mediated depletion of Evi with the *hh-Gal4* driver. Both Evi-CTD and Evi-ECD antibodies detected a strong loss of Evi staining in the posterior compartment of the wing disc when compared to the endogenous protein detected in the anterior compartment (Figure 1A-B and C-D, respectively). This data suggests that both antibodies are specific to the Evi protein.

**Figure 1.**
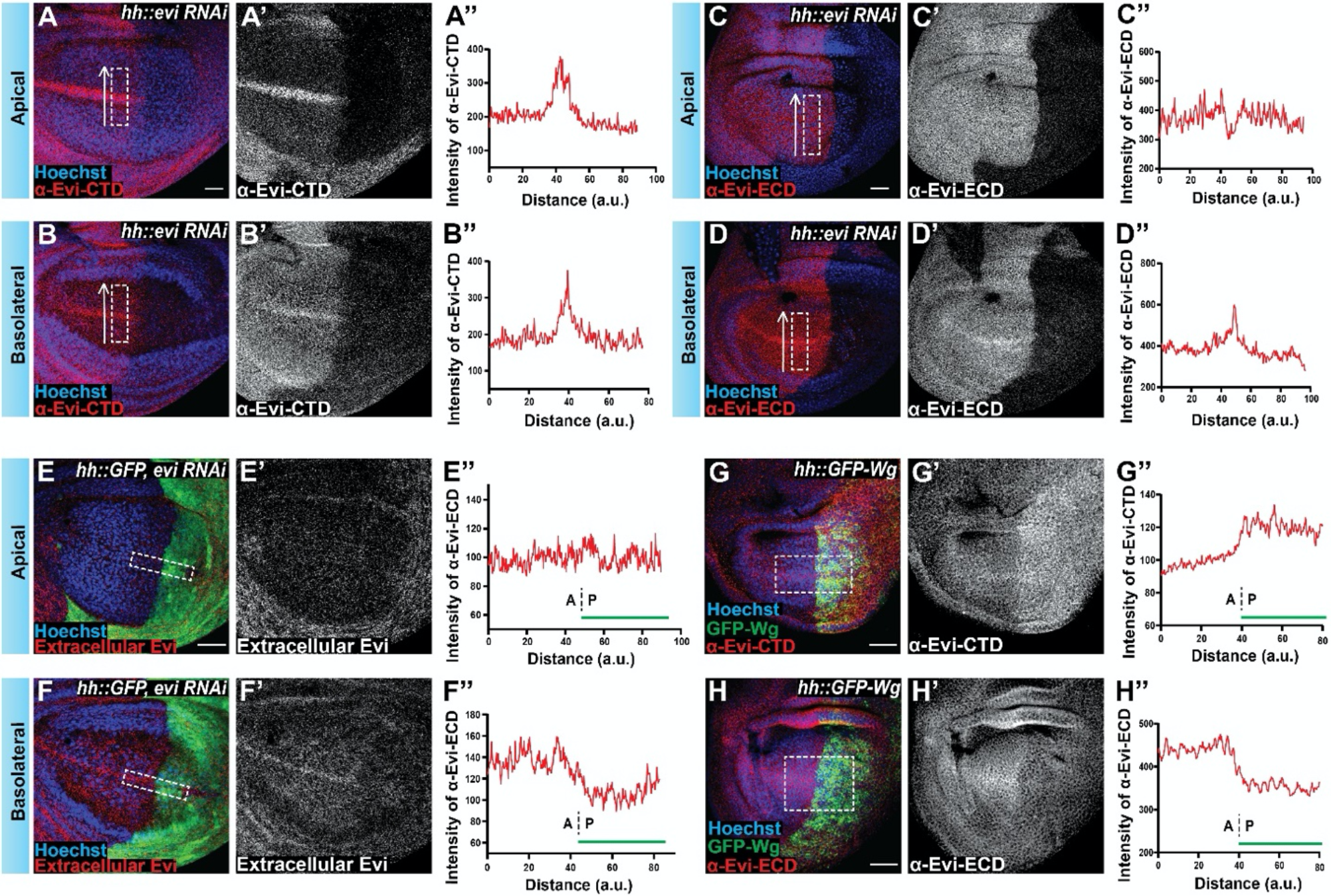
Differential labeling of Evi by α-Evi-CTD and α-Evi-ECD antibodies in the wing epithelium. **A-B)** Representative images of Evi-CTD (C-terminal domain) antibody staining (red) in the wing imaginal disc of *Drosophila* with *hh-Gal4* driven expression of *UAS-evi-RNAi* in the posterior compartment (N=4). Hoechst (blue) is used as the nuclei marker. A-A’) Shows the apical section, and A’’ shows the intensity profile of α-Evi-CTD levels across the dorsoventral (DV) boundary (box marked in A). B-B’) Shows the basolateral section and the intensity profile of α-Evi-CTD levels across the DV boundary is shown in B’’ along the box marked in B. **C-D)** Representative image showing Evi-ECD (extracellular domain) antibody staining (red) after depleting Evi protein in the posterior compartment of the wing disc (N=9). C-C’’) Shows Evi-ECD antibody staining for the apical region of the wing disc with a graph showing the intensity profile for the box marked in C. D-D’’) Images for the basolateral side of the same Evi depleted wing discs with quantification by intensity profile in D’’. **E-F)** Representative image for the extracellular Evi staining with α-Evi-ECD (red) in the wing disc expressing *UAS-evi-RNAi* using *hh-Gal4* driver in the posterior compartment, marked with GFP expressing region (green) (N=9). E-E’) Represents the apical section of the wing disc with intensity profile (along the box marked in E) in E’’. F-F’) Shows the extracellular Evi staining towards the basolateral region of the same wing disc as shown in E. F’’) Graph representing the intensity profile along the DV boundary (as marked with box in F), A and P represent anterior and posterior regions respectively for the wing disc used for quantification with a green bar at the right bottom represents RNAi expressing region. **G-H)** Representative image for the GFP-Wg overexpression (green) using a *hh-Gal4* driver for 24 hours at 29°C under controlled expression by *tub-Gal80*^*ts*^. G-G’) α-Evi-CTD staining (red) for the GFP-Wg expressing wing disc with Evi protein’s intensity profile in G’’ for the region marked with a box in G (N= 4). H-H’) Shows the α-Evi-ECD staining (red) with GFP-Wg overexpressing wing disc, and quantification for Evi protein is shown in H’’ for the box within the wing pouch marked with a white box in H (N=7). For apical and basolateral images, 2-3 confocal slices were merged. S.B.= 30μm. a.u. = arbitrary unit.

Past studies have shown that expression of Wg can stabilize Evi protein levels; thus, higher levels of Evi protein are detected in the producing cells at the DV boundary (Port et al., 2008). As expected, in the anterior control compartment of *hh-Gal4; evi-RNAi* discs, α-Evi-CTD detected strongly stabilized Evi towards the apical side of the producing cells (Figure 1A-A”), while a mild stabilization was also observed towards the basolateral side (Figure B-B”). However, when analyzed with the Evi-ECD antibody, the higher levels of Evi were not detectable at the apical side (Figure 1C-C”, compare with Figure 1A and A’’), while the basolateral Evi was still observed (Figure 1D-D”). Next, we tested the plasma membrane localization of Evi with the extracellular staining using α-Evi-ECD on *hh-Gal4; evi-RNAi* discs. Similar to the total staining, the extracellular staining detected stabilized Evi only at the basolateral side in the anterior control compartment (Figure 1F-F”), while apical stabilization was not observed (Figure 1E-E”). These results suggest that α-Evi-CTD and α-Evi-ECD detected different populations of Evi protein in the cells.

The first extracellular loop is also the binding site for Wnt (Nygaard et al., 2021; Zhong et al., 2021) thus, we hypothesized that perhaps α-Evi-ECD is unable to detect Wnt-bound Evi protein and can only identify Wnt-unbound or ‘free’ Evi. To test this, we asked if increasing the levels of Wg in the cells would show a differential effect on α-Evi-CTD and α-Evi-ECD labeling. To this end, we overexpressed GFP-Wg in the posterior compartment for 24 hours. As expected, the total Evi levels were increased upon the overexpression of GFP-Wg in the posterior compartment (Figure 1G-G”) (Port et al., 2008). However, in contrast to this, α-Evi-ECD levels were reduced (Figure 1H-H”), suggesting that α-Evi-ECD is unable to recognize Wnt-bound Evi.

### Evi remains bound to Wg while traveling from the ER to the cell surface

Higher levels of total Evi are present towards the apical side of the producing cells (Port et al., 2008) and Wg is also trafficked first to the apical surface. However, lower α-Evi-ECD labeling at the apical side indicates that the Evi may remain bound to Wg in the anterograde route to the apical surface. To test whether the Wg and Evi separation could occur in the anterograde route, we designed an assay whereby we captured and stabilized GFP-tagged Wg in the anterograde route and probed for concomitant Evi stabilization. To this end, we used the expression of an anti-GFP nanobody-based ‘morphotrap’ system directed toward the apical surface of the wing epithelium (Harmansa et al., 2017; Rothbauer et al., 2006) in the endogenously tagged *GFP-wg* background (Port et al., 2014) (Figure 2A).

**Figure 2.**
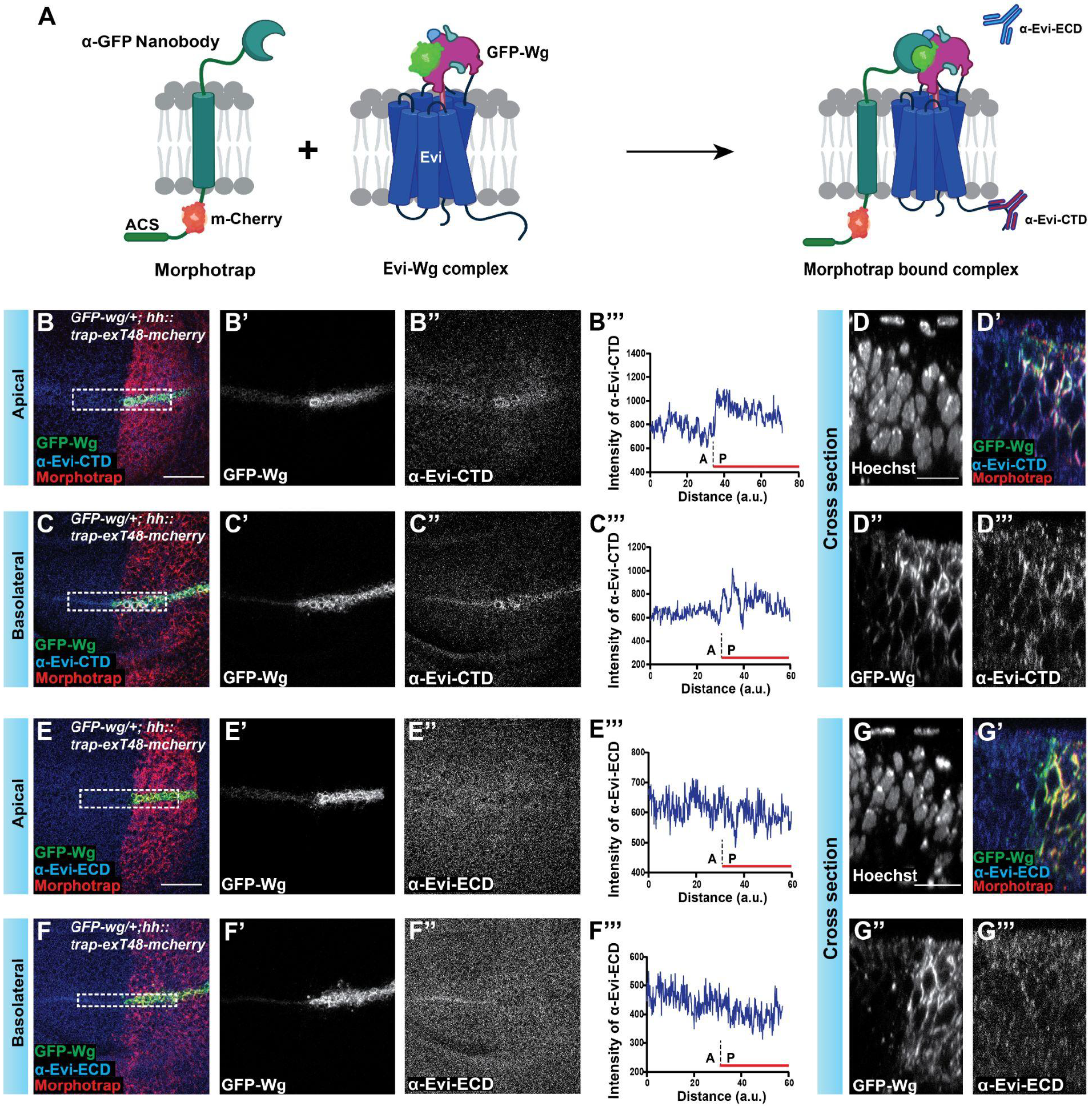
Wg and Evi travel together in the anterograde route. **A)** Schematic depiction illustrating the distinct structural domains of morphotrap and its interaction with Evi and GFP-Wg complex. **B-C)** Representative images for the apical sections (B-B’’) and basolateral (C-C’’) sections of the *GFP-wg/+* wing disc expressing morphotrap in the posterior compartment (marked red) and stained with α-Evi-CTD. Intensity profiles for Evi levels along the DV boundary are shown for the apical and basolateral side in B’’’ and C’’’, respectively; the red line marks the expression of the morphotrap. **D-D’’’)** Represents the cross section (XZ plane) across the DV boundary for the GFP-Wg accumulation and α-Evi-CTD staining; Hoechst marks nuclei (N = 5). **E-F)** Representative images for the apical (E-E’’) and basolateral sections (F-F’’) of the discs with the aforementioned genotype stained with α-Evi-ECD antibody. The intensity profile of α-Evi-ECD staining is shown for the apical and basolateral side in E’’’ and F’’’, respectively. **(G-G’’’)** Represents the cross section (XZ plane) across the DV boundary for the GFP-Wg accumulation and α-Evi-ECD staining; (N = 10). For apical and basolateral sections, 7 confocal slices were merged. S.B. = 30μm for XY plane and 10μm for XZ plane. a.u. = arbitrary unit.

We find that the expression of m-cherry tagged apical morphotrap with *hh-Gal4* in the posterior half of the homozygous *GFP-wg/GFP-wg* wing disc led to the stabilization of GFP-Wg (Figure S1A-B). However, we also observed patterning defects, which were largely visible as the duplication of the wing primordia, probably due to dysregulation of Wnt signaling activity because of mislocalization of GFP-Wg (Figure S1A and B). To avoid any possible artifacts due to the homozygous *GFP-wg* background, we expressed the apical morphotrap in the heterozygous *GFP-wg* background. Similar stabilization of GFP-Wg was observed upon the expression of the apical morphotrap in the *GFP-wg/+* wing disc without any noticeable patterning defects (Figure 2B). We further find that the stabilization of GFP-Wg was not restricted to the apical surface as it was also detected at the basolateral side (Figure 2B’, C’ and D’’), most likely due to continuous expression over a long period. More importantly, we find that the accumulation of Wg led to the concomitant increase in the total Evi levels in both *GFP-wg/GFP-wg* and *GFP-wg/+* wing discs (Figure S1A-A’’ and Figure 2B’’, C’’ and D’’’, respectively). Furthermore, in these experiments, a mild decrease in α-Evi-ECD staining was observed in the *GFP-wg/+* wing discs (Figure 2F’’ and F’’’), whereas a strong reduction was observed in *GFP-wg/GFP-wg* discs (Figure S1B-B’’), suggesting that expression of morphotrap led to capturing of GFP-Wg and this further caused most of the Evi to be trapped with GFP-Wg (Figure 2A). Altogether, these results indicate that Evi-Wg separation does not occur during the anterograde route from the ER to the plasma membrane.

### Evi and Wg are internalized together from the apical surface

Past studies in human cells have shown that Evi is internalized in a Dynamin-dependent process (Gasnereau et al., 2011). Moreover, Wg was also shown to undergo internalization through a Dynamin-dependent mechanism (Strigini and Cohen, 2000). However, whether Evi and Wg are internalized together from the apical surface of producing cells remains unknown.

To test if Wg and Evi enter the retrograde route together, we performed a pulse-chase assay for Wg and analyzed the colocalization of internalized Wg with Evi. The pulse-chase assay involved labeling of the extracellular Wg protein with a primary antibody on ice, followed by a short pulse of internalization at 30°C. The unbound antibody was removed from the extracellular space with an ice-cold acidic buffer wash, and the chase was performed for 10 mins at 25°C (Fig. 3A) (Hemalatha et al., 2016). Consistent with the previous studies, we find that the internalized Wg colocalized with the early endosomal marker Rab5 in the producing as well as receiving cells (Figure 3B–B’’ and C) (Linnemannstöns et al., 2020; Marois et al., 2006). Next, we analyzed the colocalization of Evi (with α-Evi-CTD) and Wg using adaptive deconvolution-based confocal microscopy to increase the lateral resolution. We find that the colocalization of internalized Wg with Evi was significantly higher in the Wg-producing cells as compared to the receiving cells (Figure 3D-D’’ and E). Furthermore, this colocalization was enriched towards the apical side, suggesting that Wg and Evi could be internalized together from the apical surface.

**Figure 3.**
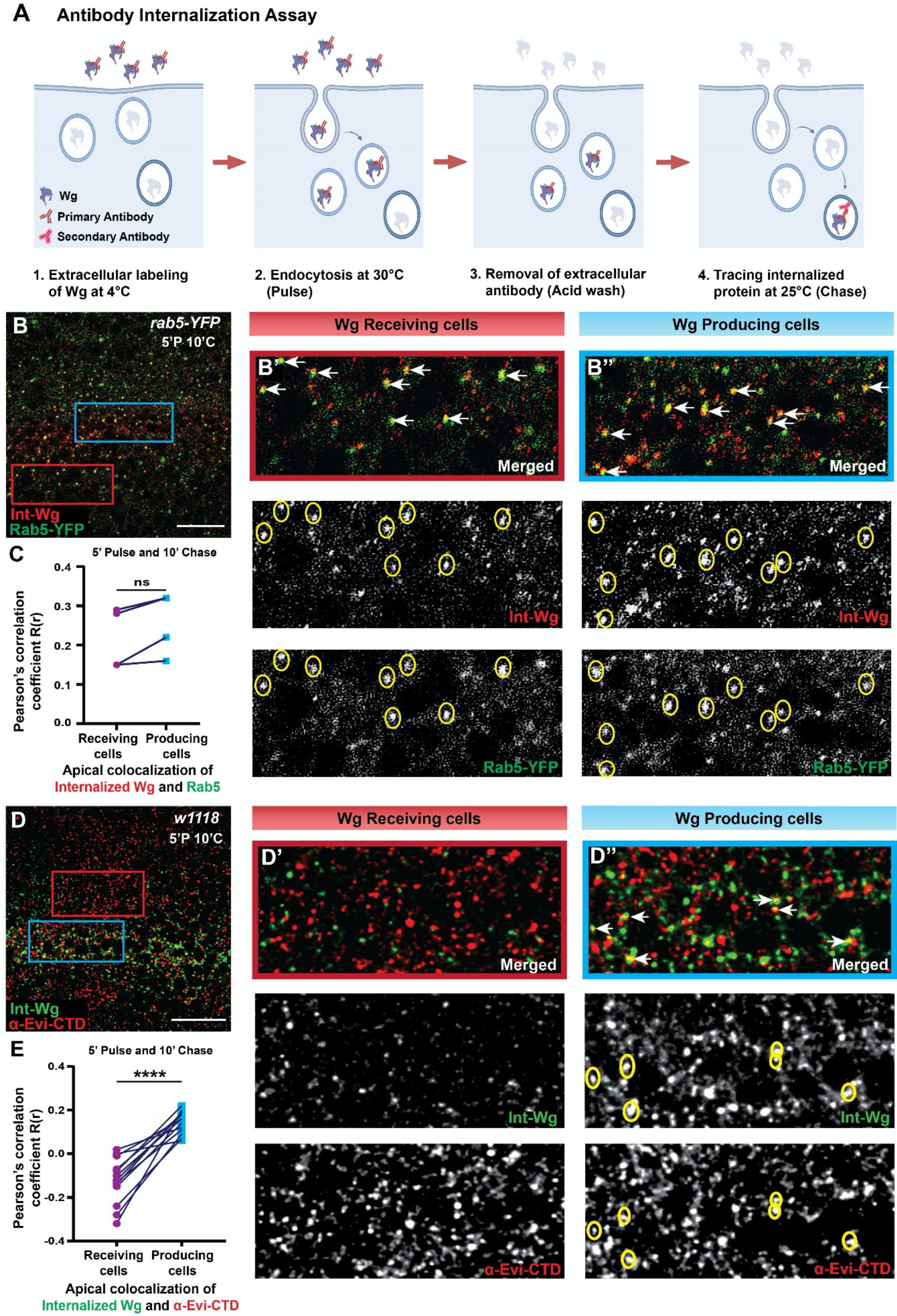
Apical internalization of Wg with Evi in the Wg-producing cells. **A)** Schematic representation of the antibody internalization assay. **B)** Representative image for the apical section (single confocal slice) of the wing epithelium with endogenously YFP-tagged Rab5 (green) and internalized Wg (red) with 5 min pulse and 10 min chase. B’ and B’’ shows the enlarged view of the colocalization of YFP-Rab5 with internalized Wg in the Wg-receiving cells (marked with red box in B) and Wg-producing cells (marked with blue box in B), respectively. **C)** Graph representing the quantification for colocalization between Wg and YFP-Rab5 using Pearson’s correlation coefficient R(r) for Wg-receiving cells (B’) and Wg-producing cells (B’’) towards the apical region of the wing disc (N = 4). **D)** Representative images of a single apical slice of the wildtype discs with internalized Wg (5’P and 10’C) (green) and α-Evi-CTD labeling (red). Colocalization of internalized Wg and total Evi in the Wg-receiving cells is shown in D’ and in the Wg-producing cells is shown in D’’. **E)** Graph showing colocalization between Evi and internalized Wg in Wg-receiving cells and Wg-producing cells towards the apical region of the wing disc, using Pearson’s correlation coefficient R(r) (N = 13). S.B = 15μm.

To further strengthen these conclusions, we performed experiments to transiently enhance early endosomal activity by expressing constitutively active YFP-tagged Rab5 (Rab5^CA^-YFP) (Stenmark et al., 1994). To control the expression of Rab5^CA^-YFP, we used a temperature-sensitive Gal80, driven under a tubulin promoter, which allows temporal regulation of Gal4 activity. *hh-Gal4, UAS-Rab5*^*CA*^*-YFP* larvae were reared until the third-instar stage at 18°C (to block Gal4 activity). Larvae were then shifted to 29°C (permissive for Gal4 activity) for 6 hours to allow the expression of Rab5^CA^-YFP. The cells expressing Rab5^CA^-YFP showed enlarged endosomes, consistent with previous observations (Stenmark et al., 1994). Wg chased for 10 mins after internalization showed strong accumulation in the enlarged endosome (Figure S2A-A2). More importantly, strong colocalization of α-Evi-CTD with Wg in Rab5^CA^-YFP compartments was observed specifically in the producing cells (Figure S2A1 and S2B), confirming that Wg and Evi are internalized together.

Next, we analyzed if Wg and Evi separation occurred in these enlarged early endosomes. The α-Evi-ECD staining showed that ‘free’ Evi levels were unchanged in the posterior compartment by the expression of Rab5^CA^-YFP (Figure S2C-C2 and S2D). Furthermore, free Evi did not significantly colocalize with Rab5^CA^-YFP, suggesting that Wg remained bound to Evi in these endosomes.

### Wg and Evi separation occurs in the acidic late endosomes

Evi has been shown to colocalize with the retromer positive late endosomes, and it is then trafficked to Golgi in a retromer-dependent manner (Belenkaya et al., 2008; Franch-Marro et al., 2008) but whether these acidic late endosomes is the site for separation of Evi and Wg is not known. We next tested if the internalized Wg is also trafficked up to late endosomes in the Wg-producing cells. RFP-tagged Vps35 was used for labeling the retromer positive endosomes, and a strong colocalization was observed with internalized Wg (Figure 4A-A’’’). However, the colocalization of free Evi with RFP-tagged Vps35 was only mildly higher in the producing cells as compared to the receiving cells (Figure 4B-B’’’). This is probably because free Evi remains only transiently in the retromer positive endosomes due to recycling.

**Figure 4.**
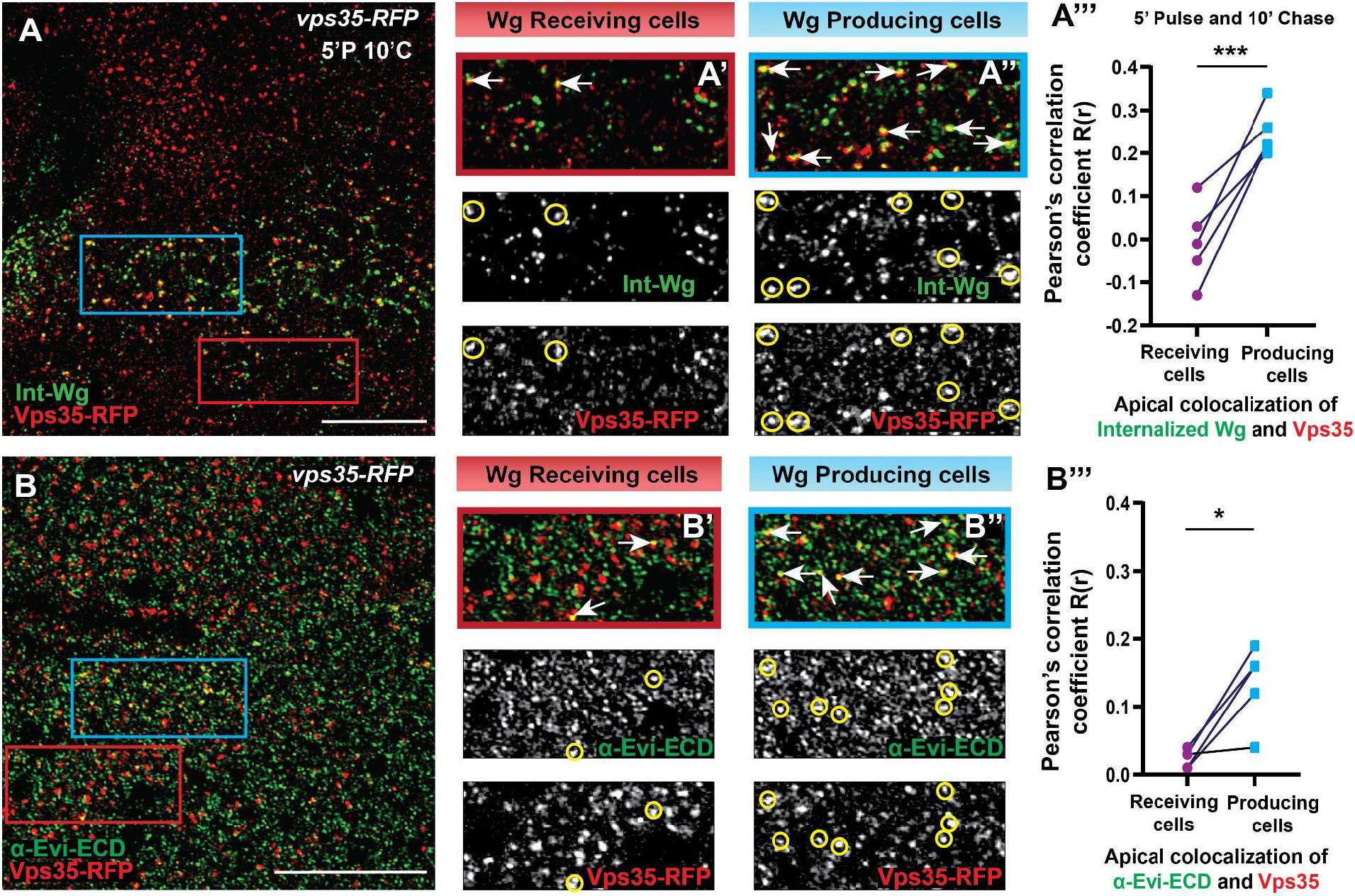
Internalized Wg reaches up to retromer positive endosomes. **A)** Representative image displaying the wing imaginal disc, depicting internalized Wg (green) subsequent to 5 min pulse (5’P) and 10 min chase (10’C) assay, along with Vps35-RFP-labeled endosomes (red). An enlarged view of the colocalization of internalized Wg (5’P 10’C) and Vps35-RFP is shown in A’ for the receiving cell (red box in A) and in A’’ for the producing cells (blue box in A). **A’’’)** Graph representing the quantification using Pearson’s correlation coefficient R(r) for the colocalization in Wg-producing as well as receiving cells for internalized Wg and Vps35-RFP towards the apical region of wing disc (N = 5). **B)** Representative image for the α-Evi-ECD and Vps35-RFP colocalization, the enlarged view for Wg-producing cells (marked with blue box) is shown in B’’ and for Wg-receiving cells (marked with red box) is shown in B’. **B’’’)** Graph represents the colocalization between α-Evi-ECD and Vps35-RFP for receiving as well as producing cells towards the apical region of the wing disc. Single slice towards the apical region was used to represent colocalization. S.B = 15μm.

Next, we checked if increasing vesicular acidification would enhance Evi and Wg separation and, therefore, the detection of ‘free’ Evi. To this end, we choose to downregulate Vps34, which is a PI3P kinase protein crucial for late endosomal fusion with lysosomes, and loss of Vps34 was shown to enhance Rab7 activity and create enlarged late endosomes (Jaber et al., 2016; Johnson et al., 2006), that are acidic in nature (Ronan et al., 2014). Thus, we expressed *vps34* RNAi in the posterior compartment of the wing discs. As expected, depletion of Vps34 showed a significant increase in the lysotracker-stained acidic vesicles localized mostly towards the apical region (Figure S3A-B’). Furthermore, the depletion of Vps34 also showed a strong increase in the levels of late endosomal marker Rab7 (Figure 5A-A’, Figure S4A’ and Figure S4C’).

**Figure 5.**
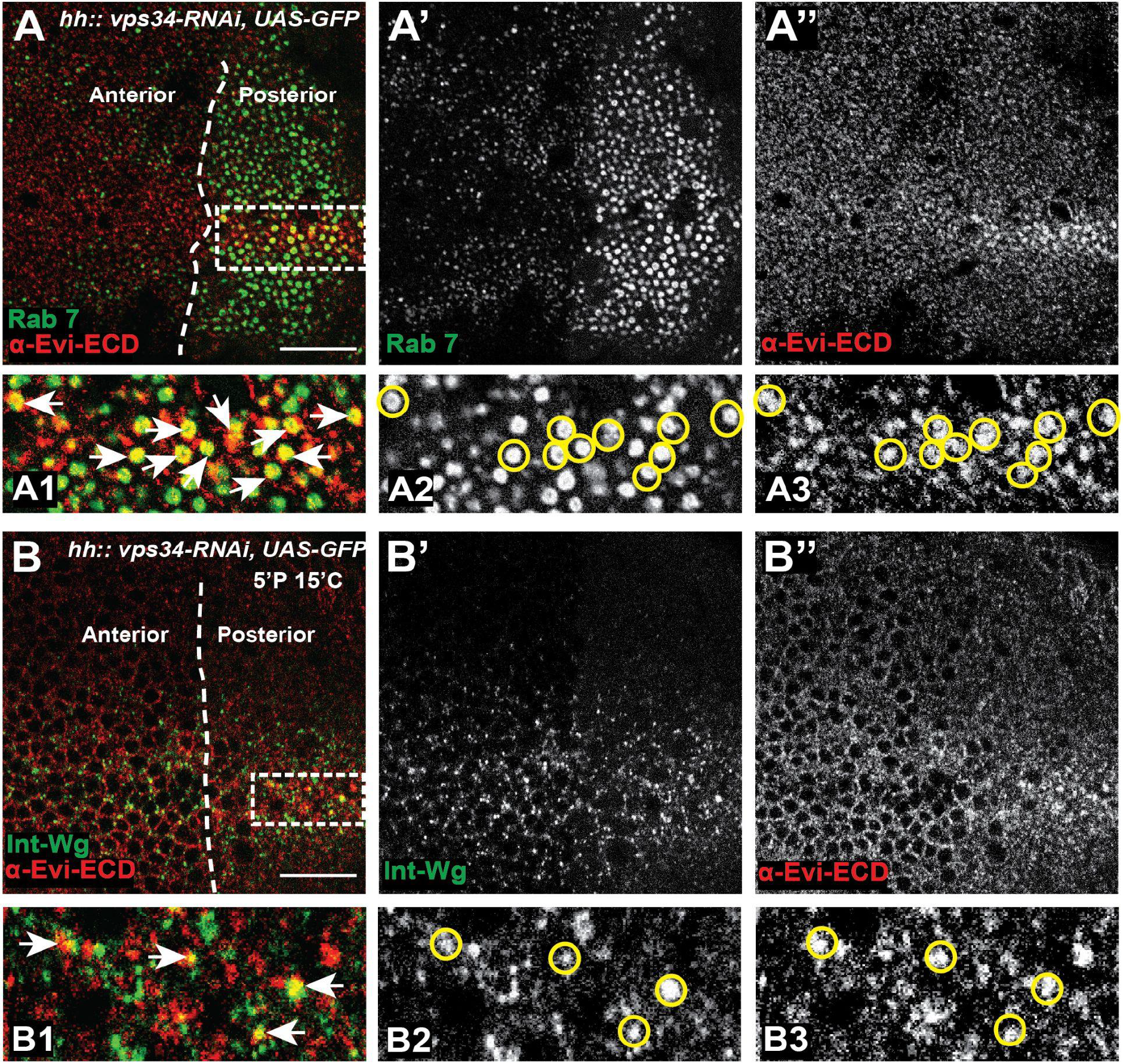
Evi-Wg complex dissociates in the late endosomes. **A)** Representative images of the apical section of the wing disc expressing *vps34*-RNAi with *hh-Gal4*, stained with Rab7 (A’, green) and α-Evi-ECD (A’’, red). A1-A3) Enlarged images of the DV boundary region in the posterior compartment marked with the white box shows colocalization between Rab7 (A2) and α-Evi-ECD (A3) (N=9). **B)** Representative images of the apical section of the wing discs with the same genotype as in A, stained for internalized Wg (5’P 15’C) and α-Evi-ECD. B1-B3) Enlarged images of region marked in B (white box) showing colocalization of internalized Wg (B2) and α-Evi-ECD (B3); (N = 6). Single slice towards the apical region was used to represent colocalization in A and B. S.B = 15μm.

With the α-Evi-CTD staining, we observed that Evi protein exhibited an accumulation within the expanded late endosomes characterized by Rab7 markers, specifically towards the apical region of the wing disc (Figure S4A-A’’ and Figure S4C-C’’). Conversely, Evi protein levels decreased towards the basolateral side of polarized cells (Figure S4B-B’’ and Figure S4C-C’’). More importantly, higher levels of free Evi were visible towards the apical side, which colocalized with Rab7 (Figure 5A-A3). Furthermore, both internalized Wg as well as total Wg showed colocalization with α-Evi-ECD positive vesicles (Figure 5B-B3 and Figure S4D-E’’ respectively). These results indicate that the loss of Vps34 enhanced the dissociation of Evi and Wg in the enlarged late endosomes. Altogether, this suggests that the Evi-Wg complex enters into the retrograde route and reaches up to the late endosomes, where they dissociate due to the acidic environment.

## DISCUSSION

Wnt molecules are lipid-modified, and therefore, they are hydrophobic in nature. Studies have shown that for the dispersion in the extracellular space, Wnt proteins bind with various carrier molecules, for example, Lipoprotein particles, Secreted Wingless Interacting Molecule (SWIM), and Heparan Sulfate Proteoglycans (HSPGs). Moreover, they can be packaged on MVB-derived exosomes and are also shown to be released on actin-rich extensions called Cytonemes (Gradilla et al., 2018; Mehta et al., 2021). Interestingly, these carriers were shown to participate in the secretion of *Drosophila* Wg protein from the polarized producing cells in the developing larval wing epithelium. However, the intracellular mechanisms required to direct the Wg proteins to different carriers are not well understood. It is believed that the internalization of Wg from the cell surface will facilitate its packaging on its carriers like exosomes and lipoproteins (Gross et al., 2012; Panáková et al., 2005).

In the case of the polarized epithelial cells, Wg is first trafficked to the apical surface, and it is then internalized for re-secretion via both apical and basolateral recycling routes. Moreover, the Wnt-cargo receptor Evi, which is essential for Wg secretion, is localized predominantly towards the apical side. Whether Wg dissociates from Evi in the anterograde route and further moves in an Evi-independent manner is not understood. In this *in vivo* study, we found that Evi and Wg separation does not occur in the anterograde route from Golgi to the plasma membrane, and Wnt-bound Evi reaches the apical membrane of the polarized-producing cells. Furthermore, our results show that Wg and Evi are internalized together and reach up to late endosomes that have a higher acidic lumen.

Our Evi antibody against the ECD specifically detected free Evi, which was observed towards the basolateral surface of the cells, while total Evi resided more strongly towards the apical side. Comparison of the free Evi with the total Evi allowed us to analyze the process of Wg and Evi separation. Vacuolar acidification has been suggested to mediate the separation of Wnt and Evi in cultured human cells (Coombs et al., 2010). In agreement with this, we find that α-Evi-ECD levels were increased upon increasing endosomal acidification by the depletion of Vps34 (Figure 5 A-A’’).

Our results are consistent with the finding that internalization is required for Wg secretion (Strigini and Cohen, 2000). However, the precise understanding of further channeling of the dissociated Wg from the late endosomes to the various polarized secondary pathways remains unclear. Studies have shown that Wg can be recycled back to the apical membrane in a Ykt6 and Rab4-dependent manner (Gross et al., 2012; Linnemannstöns et al., 2020). Besides, Wg can also undergo transcytosis from the apical to the basolateral side in a Godzilla-dependent manner (Yamazaki et al., 2016). Interestingly, the Ykt6-Rab4-mediated recycling was suggested to operate at the early endosomal levels and was shown to be independent of Evi recycling. Moreover, Godzilla is also localized at the early endosomes and regulates recycling via the ubiquitination of VAMP3 (Yamazaki et al., 2013). It will be further interesting to analyze the effect of these pathways on the separation of the Evi-Wg complex.

Polarized trafficking of Wnt proteins is also observed in the vertebrate model system. Studies using polarized Madin–Darby canine kidney (MDCK) cells have shown that Wnt3a and Wnt11 follow separate polarized secretion routes beyond Golgi due to their different pattern of glycosylation. While Wnt3a was shown to be trafficked to the basolateral side in Evi/Wls dependent process, Wnt11 was released from the apical side in Evi/Wls independent manner (Yamamoto et al., 2013; Yamamoto et al., 2015). Interestingly, the Wg-expressing cells at the DV boundary also express the Wnt6 ligand (Gracia-Latorre et al., 2022; Janson et al., 2001). However, only a single Evi is present in *Drosophila*, which would mediate the secretion of both Wg and Wnt6. Whether these two ligands follow similar polarized secretion routes remains to be seen.

## MATERIALS AND METHODS

### *Drosophila* Stocks

Knockdown by RNAi and overexpression of particular genes were done using upstream activation sequence (UAS) fly lines as follows: *UAS-evi-RNAi* KK (VDRC 103812), *UAS-vps34-RNAi* KK (VDRC 100296), *LexA-UAS-trapExT48-mcherry* (BDSC 68177), *UAS-YFP-rab5 Q88L* (BDSC 9774), *UAS-GFP (*3rd chromosome), *UAS-GFP-wg (Pfeiffer et al*., *2002)*. To express UAS constructs, we used *hh-Gal4* (Tanimoto et al., 2000) and *tub-Gal80*^*ts*^ (BL7108) to control expression temporally. Other fly lines used were: *rab5-EYFP* (BDSC 62543), *vps35-RFP* (BDSC 66527), *GFP-wg* ((Port et al., 2014)), and *w1118* (BDSC 3605).

### Antibodies

Larval wing imaginal discs were stained using the following antibodies: mouse anti-Wg (1:50 for total staining) Developmental Studies Hybridoma Bank (DSHB), mouse anti-Rab7 (1:10) (DSHB), rabbit α-Evi-CTD (1:100) (generated for this study; based on Port et al., 2008) and rabbit α-Evi-ECD (1:100) (generated for this study). Secondary antibodies used for fluorescent labeling were Alexa-405, Alexa-488, Alexa-594, Alexa-647 (Invitrogen) at 1:500 dilutions and Hoechst 33342, H3570 (1:1000, Invitrogen).

### Immunostaining

*Drosophila* third instar larvae were dissected and stained using the standard immunostaining protocol. The larval head complexes with wing discs were dissected in 1X PBS and fixed with 4% paraformaldehyde for 30-40 minutes at room temperature. Subsequently, the samples were treated with PBS-T (0.2% Triton in PBS), blocked using BBT (0.1% BSA in PBS-T), and then exposed to the primary antibody overnight at 4°C with gentle agitation. Following multiple washes with PBS-T, fluorescent dye-labeled secondary antibodies were used for 1-2 hrs incubation at room temperature, afterwards PBS-T washes were performed before mounting the tissue on slides having Vectashield (Vector Labs) as mounting media. The transparent tape was applied on the glass slides prior to tissue mounting to achieve precise elevation, facilitating optimal tissue preservation within the mounting media.

For extracellular staining of Evi protein, the wing discs containing head complexes were incubated with primary antibody, i.e., α-Evi-ECD, with 1:25 dilution in blocking (2% BSA in PBS-T), overnight at 4°C after fixation, then proceeded conventionally for fluorescently labeled secondary antibody staining and mounting.

Lysotracker Deep red (Invitrogen L12492) staining was done with 1:2000 dilution in Schneider’s media, and wing discs containing larval heads were incubated in it for 10 min at 30°C. After rinsing the head complexes with PBS, the wing discs were mounted in Vectashield for confocal imaging.

### Antibody Internalization Assay

The antibody internalization assay was performed as mentioned in Hemlatha et al., 2016. In summary, wandering third instar larvae were dissected in Schneider’s media, the wing imaginal discs were incubated on ice for 5 min. Wing discs were further incubated with Wg antibody (1:25 dilution in Schneider’s media) for 30 min at 4°C. Pulse was given at 30°C with fresh Schneider’s media for 5 min. The acid stripping of extracellular antibodies was done with ice-cold 0.1M Glycine (pH 3.5) for 30-40 sec before proceeding for the time-point dependent chase at 25°C time points. Before mounting, standard staining protocols were followed to fix and stain the sample with other antibodies.

### Antibody Generation and Purification

A 174 amino acids long extracellular loop 1 (from amino acid position 66 to 238) of *Drosophila* Evi protein was chosen for antibody production. The corresponding 522bp long DNA fragment was amplified with HindIII and XhoI sites and cloned in an IPTG inducible pET28a(+) vector with a C-terminal 6X histidine tag. The protein was expressed in *E. coli* bacteria at 18°C. The recombinant protein was purified using the Ni-NTA (Biorad IMAC Ni-Charged Resins) affinity chromatography under denaturing conditions. The purified recombinant protein was further dialyzed and concentrated using the membrane columns (Pierce Protein Concentrator 3KMWCO). This purified protein was injected as an antigen in two rabbits. The serum was collected 10 weeks after injection. The Evi antibody was purified from the serum via western blotting (Fang, 2012). After incubating the blot with serum, the antibodies bound to Evi (extracellular loop) on the nitrocellulose membrane were isolated through elution with pH 2.5 Glycine solution. These eluted antibodies were subsequently neutralized with Tris buffer (pH 8.0) and used as α-Evi ECD staining in our experiments.

### Microscopy and Image Acquisition

Immunostained wing discs were imaged with an Olympus FV3000 live cell microscope with approximately 1K resolution (40X, 60X, and 100X oil objective lens). The image represented in Figure 3D was captured using an adaptive deconvolution-based lightning Leica confocal microscope (63X oil objective lens) to obtain high-resolution images (approximately 2K). The images were acquired with 1μm step size to obtain a z-stack for a sample with 1X and 3X optical zoom. The cross-section was taken with a line scan having 0.05μm step size to form an XZ plane.

### Image Processing

After acquiring the images using a confocal microscope, the images were processed in ImageJ (Fiji) and Adobe Photoshop CS6 v13.0. Image quality and signal resolution were enhanced post-acquisition by the deconvolution of images in the CellSens software of the Olympus microscope. Figures were made in Adobe Illustrator (Adobe Illustrator CS6 Tryout version 16.0.0), and models were created with BioRender.com.

### Colocalization and Statistical Analysis

Colocalization analyses were performed with the help of CellSens software. ROIs from the DV boundary and outside regions were used for colocalization analysis in the wing pouch region. By applying a specific threshold value for each channel, only puncta having appropriate intensity were considered for colocalization, and Pearson correlation coefficient R(r) values were obtained for each plane separately. Single apical confocal slices (3-4) were used to obtain the average value of Pearson’s correlation coefficient R(r). The data was analyzed using Microsoft Excel, and Graphs were plotted using GraphPad Prism version 8.0.2. Each data point in the graph for colocalization represents Pearson’s value R(r) between two channels, and the lines connecting the data points represent the same sample for comparing the control (the region outside the DV boundary) and experimental region (DV boundary). For statistical analysis, paired t-tests were used to determine the significance level between the control and experiment. For the intensity plot, the images were quantified in ImageJ by plot profile, and these values were plotted in the graph by GraphPad Prism.

## AUTHOR CONTRIBUTIONS

Conceptualization: Satyam Sharma and Varun Chaudhary; investigation: Satyam Sharma and Varun Chaudhary; formal analysis: Satyam Sharma and Varun Chaudhary; methodology: Satyam Sharma and Varun Chaudhary; Validation: Satyam Sharma and Varun Chaudhary; writing—original draft preparation: Satyam Sharma and Varun Chaudhary; writing—review and editing: Satyam Sharma and Varun Chaudhary; supervision: Varun Chaudhary; project administration: Varun Chaudhary; funding acquisition: Varun Chaudhary. All authors have read and agreed to the published version of the manuscript.

## ACKNOWLEDGMENTS

We are thankful to V.C. laboratory members and Dr. Sunando Datta (IISERB) for the helpful discussion on the project and manuscript. We thank Dr. Vimlesh Kumar (IISERB), BDSC, and VDRC for *Drosophila* stocks and reagents. We are also grateful for the IISER Bhopal Fly facility and the DST-FIST facility for the confocal microscopy.

## DATA AVAILABILITY STATEMENT

The data that supports the findings of this study are available in the supplementary material of this article.

