## Supplemental data for "*Drosophila* Wg and Evi/Wntless dissociation occurs post apical internalization in the late endosomes"

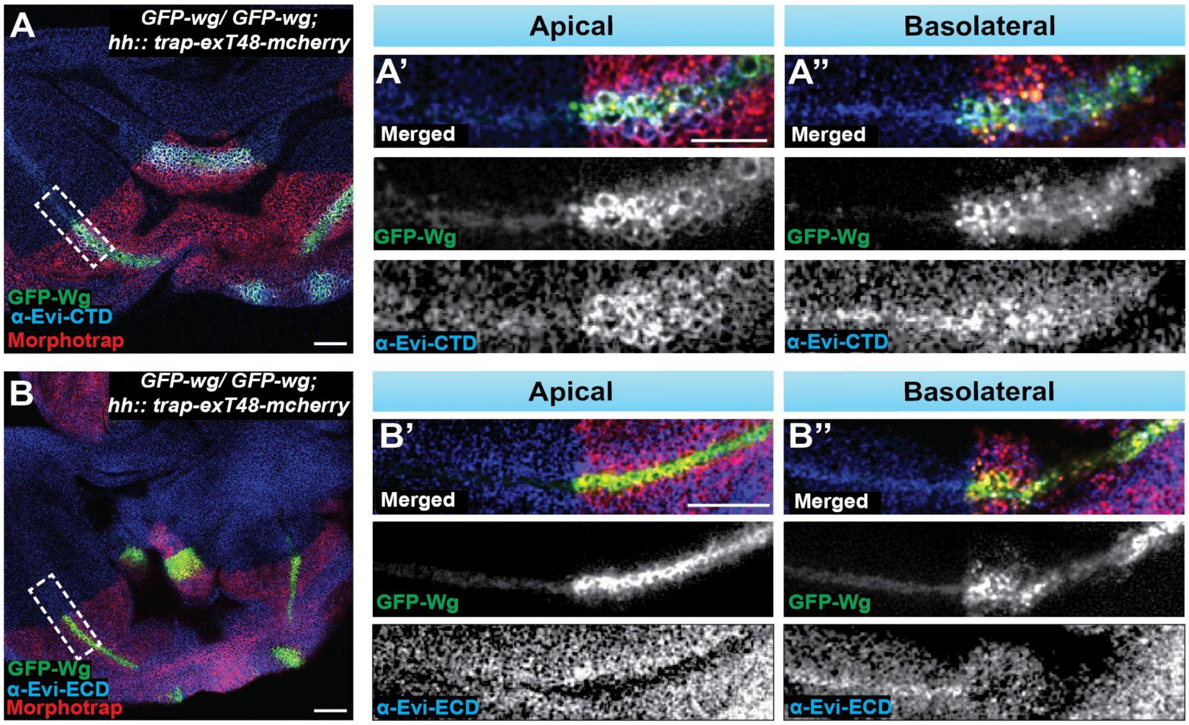


**Figure S1 (Related to Figure 2) - GFP-Wg-bound Evi levels increased by the expression of morphotrap on the homozygous *GFP-wg discs.***

**A-B)** Representative images showing *GFP-wg* homozygous discs expressing the morphotrap in the posterior compartment with *hh-Gal4* stained with either α-Evi-CTD (A-A'') or α-Evi-ECD (B-B''). A' shows the apical levels of α-Evi-CTD staining and GFP-Wg stabilization. A'' shows the basolateral levels of α-Evi-CTD and GFP-Wg; (N = 9). B' shows the apical levels of α-Evi-ECD staining and GFP-Wg stabilization. B'' shows the basolateral levels of α-Evi-ECD and GFP-Wg; (N = 12). 4 confocal slices were merged for apical and basolateral images. S.B.= 30μm.

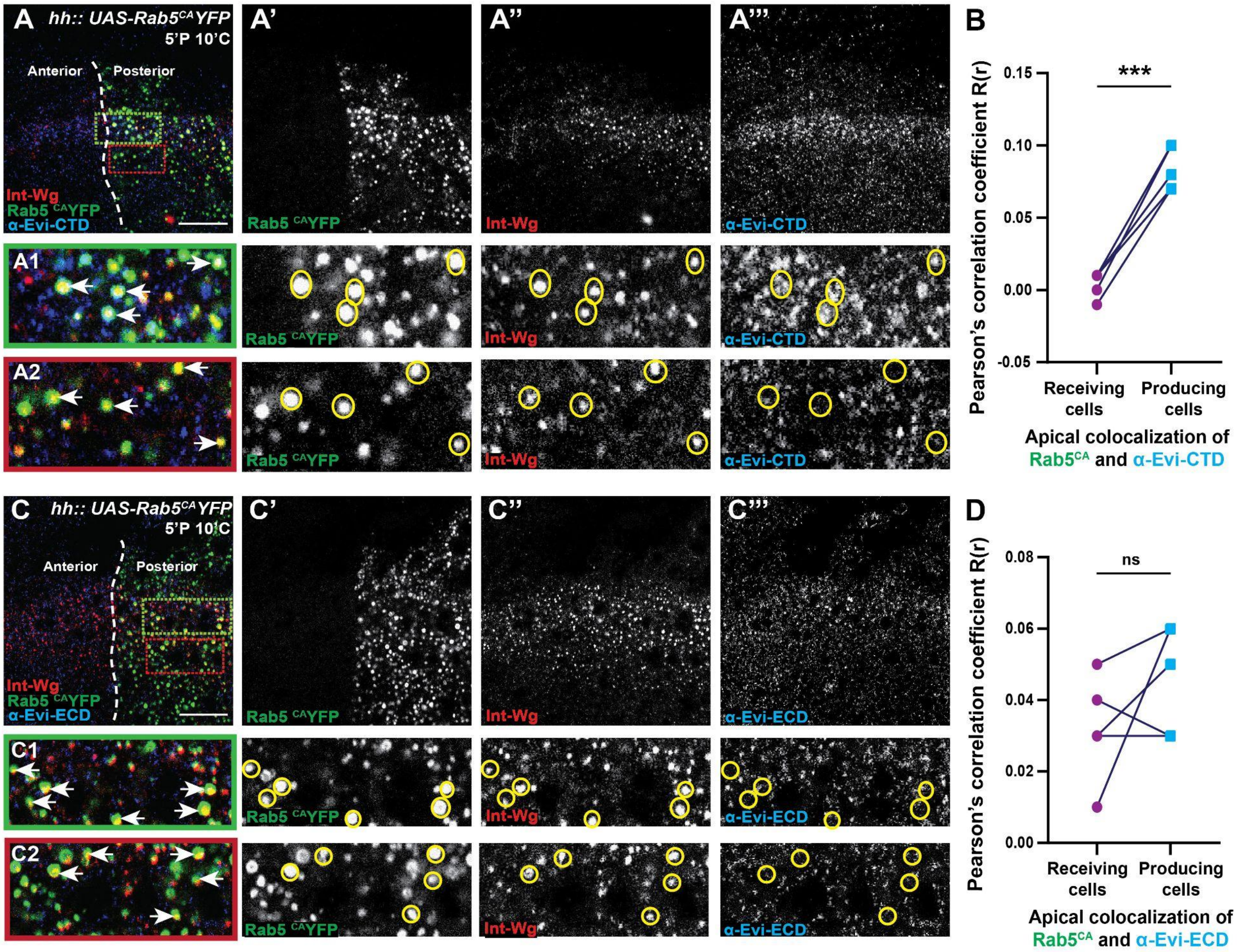


**Figure S2 (Related to Figure 3) - Early endosomal localization of Evi-Wg complex**.

**A-D)** Representative image showing the apical section of the wing imaginal disc expressing Rab5$^{CA}$-YFP (A' and C', green) using *hh-Gal4* in the posterior compartment with internalized Wg (5'P 10'C) (A'' and C'', red) and Evi labeling with either α-Evi-CTD (A''', in blue) or α-Evi-ECD (C''', blue). A1 and A2 show an enlarged view of the colocalization between Rab5$^{CA}$-YFP, Evi-CTD labeled puncta, and internalized Wg in the producing cells at the DV boundary (A1, green box marked in A) and in the receiving cells (A2, red box in A) (N = 5). An enlarged view of the colocalization between α-Evi-ECD, Int Wg, and Rab5$^{CA}$ along the DV boundary is shown in **C1** and in receiving cells in **C2** (N = 5). Colocalization analysis between Rab5$^{CA}$-YFP and Evi protein is shown as a graph for α-Evi-CTD in **B** and α-Evi ECD in **D,** for the apical region of the wing disc. Single slice towards the apical region was used to represent colocalization in A and C. S.B = 15μm**.**

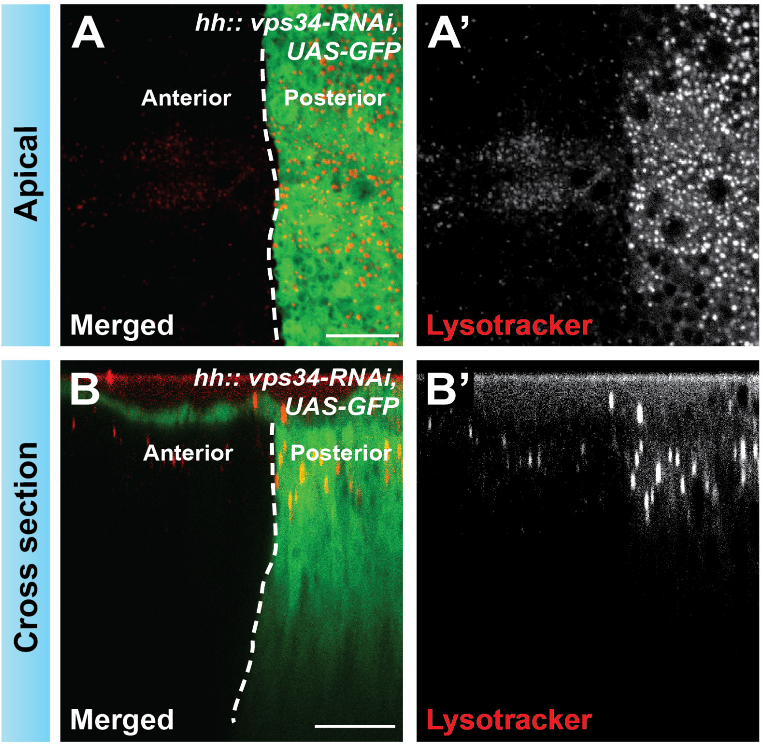


**Figure S3 (Related to Figure 5) - Loss of Vps34 increases acidic vesicles apically.**

**A)** Representative images for the apical region of the wing disc expressing *vps34*-RNAi in the posterior compartment using *hh-Gal4* driver (marked by GFP in green), acidic vesicles are stained with Lysotracker (A', red). **B-B')** Cross-sectional (XZ) view for the same wing disc along the DV boundary representing lysotracker staining (N =6). 4 slices were merged towards the apical region of the wing discs to represent the apical section in A. S.B. = 15 μm for XY and XZ planes.

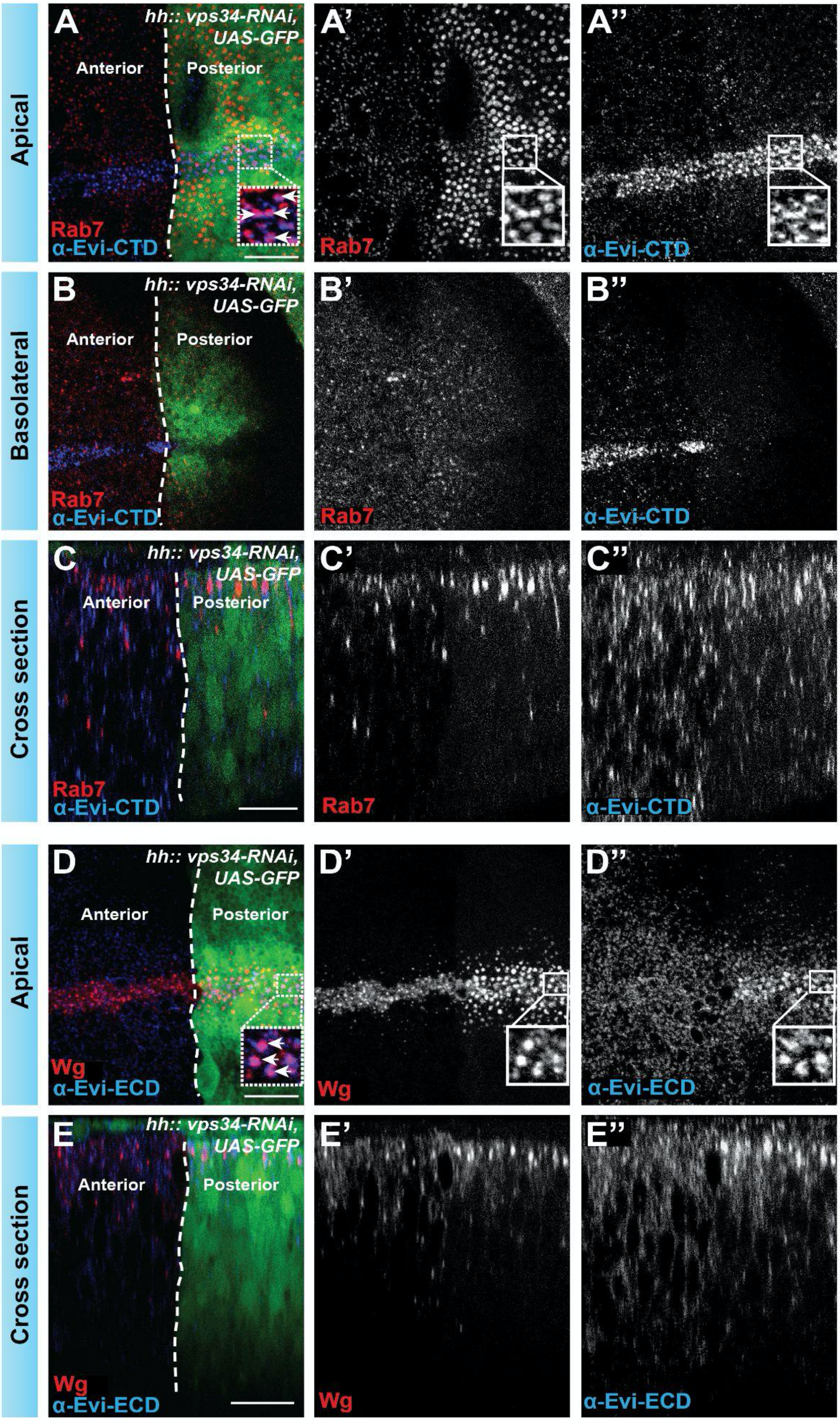
Apical
A
hh:: vps34-RNAi, UAS-GFP
Anterior
Posterior
Rab7
α-Evi-CTD
A'
Rab7
A''
α-Evi-CTD
Basolateral
B
hh:: vps34-RNAi, UAS-GFP
Anterior
Posterior
Rab7
α-Evi-CTD
B'
Rab7
B''
α-Evi-CTD
Cross section
C
hh:: vps34-RNAi, UAS-GFP
Anterior
Posterior
Rab7
α-Evi-CTD
C'
Rab7
C''
α-Evi-CTD
Apical
D
hh:: vps34-RNAi, UAS-GFP
Anterior
Posterior
Wg
α-Evi-ECD
D'
Wg
D''
α-Evi-ECD
Cross section
E
hh:: vps34-RNAi, UAS-GFP
Anterior
Posterior
Wg
α-Evi-ECD
E'
Wg
E''
α-Evi-ECD

FigureS4

**Figure S4 (Related to Figure 5) - Loss of Vps34 leads to Evi accumulation in the late endosomes and decreased basolateral levels.**

**A-B)** Representative image for the apical section (A-A'', merge of 4 slices) and basolateral section (B-B'', merge of 4 slices) of the wing disc expressing *vps34* RNAi in the posterior half using *hh-Gal4; UAS-GFP* (green), with Rab7 (A' and B', red) and α-Evi-CTD (A'' and B'', blue) antibody stainings. The white boxes represent the enlarged view (as insets) of the apical colocalization between Rab7 and α-Evi-CTD. **C-C'')** A cross-sectional view of the wing disc along the DV boundary from the disc shown in A (N = 4)**. D)** Representative image for Wg (D', in red) and α-Evi-ECD (D'', blue) staining using the same genotype as shown in A with insets representing the colocalizing puncta between Wg and Evi protein (merge of 4 slices). **E-E'')** A cross-sectional view along the DV boundary for the disc is shown in D (N = 4)**.** S.B. = 15μm for XY and XZ planes.
